# Data management and sharing in neuroimaging: Practices and perceptions of MRI researchers

**DOI:** 10.1101/266627

**Authors:** John A. Borghi, Ana E. Van Gulick

**Author notes:** these authors contributed equally to this work.

## Abstract

Neuroimaging methods such as magnetic resonance imaging (MRI) involve complex data collection and analysis protocols, which necessitate the establishment of good research data management (RDM). Despite efforts within the field to address issues related to rigor and reproducibility, information about the RDM-related practices and perceptions of neuroimaging researchers remains largely anecdotal. To inform such efforts, we conducted an online survey of active MRI researchers that covered a range of RDM-related topics. Survey questions addressed the type(s) of data collected, tools used for data storage, organization, and analysis, and the degree to which practices are defined and standardized within a research group. Our results demonstrate that neuroimaging data is acquired in multifarious forms, transformed and analyzed using a wide variety of software tools, and that RDM practices and perceptions vary considerably both within and between research groups, with trainees reporting less consistency than faculty. Ratings of the maturity of RDM practices from ad-hoc to refined were relatively high during the data collection and analysis phases of a project and significantly lower during the data sharing phase. Perceptions of emerging practices including open access publishing and preregistration were largely positive, but demonstrated little adoption into current practice.

## Introduction

Magnetic resonance imaging (MRI) is a popular and powerful neuroimaging technique for investigating the structure and function of the human brain. Functional MRI (fMRI), which enables researchers to assess activity in specific brain areas over time by measuring changes in blood oxygenation^1^, has been particularly influential in clinical and cognitive neuroscience^2^. Like their peers in social psychology^3^ and other data-intensive disciplines^4^, neuroimaging researchers have grappled with questions related to the rigor and reproducibility of their methods. As a result, there has been a substantial amount of discussion within the field about the need to foster open science practices including the regular sharing and reuse of research data^5^. However, it is unclear to what extent such practices have been adopted by the active research community.

Outside the laboratory, research data has also become an increasing focus for academic libraries. Though issues of rigor and reproducibility have been explicitly addressed in some library activities^6^, the majority of data-related library services are focused on research data management (RDM). Providing a single comprehensive definition of RDM is difficult due to the number of stakeholders involved, but the term generally encompasses topics related to how data and other research materials are documented, curated, and preserved^7^. Issues related to research design and analysis (e.g. power, preprocessing and analysis methods) are generally not considered part of RDM and effective RDM practices are not necessarily synonymous with those associated with open science as data can be well managed but not made openly available. However, effective RDM is crucial to establishing the accessibility of data and other research materials after a project’s conclusion and openly available data are only useful if saved, documented, and organized, in a manner that enables examination, evaluation, and reuse by others.

Though the configuration of services varies considerably between institutions, library RDM initiatives generally emphasize skills training and assisting researchers in complying with data-related policies and mandates^8^. In this role, academic librarians, some with extensive research backgrounds in addition to their information science training, are able to contribute their expertise to active research projects. Major challenges for library RDM initiatives include the degree to which RDM-related practices and perceptions vary between research disciplines^9^ and change over time^10^. Potentially significant differences between researchers and librarians in their perceptions and priorities surrounding data have been less explored, but likely also represent substantial challenges. As a highly interdisciplinary field currently grappling with issues closely related to RDM, neuroimaging research involvin MRI represents an ideal case study for thoroughly examining how active researchers are currently managing and sharing their data.

The complexities inherent in collecting, analyzing, and disseminating MRI data underscore the necessity of establishing well-defined RDM practices in neuroimaging research. Even a relatively straightforward project involving MRI requires the management of data in a variety of forms from a variety of sources. In addition to the data collected using an MRI scanner, this may include sensitive medical information (e.g. pregnancy status, psychiatric and medical diagnoses), task-related behavioral data (e.g. response accuracy, reaction time), and questionnaire responses. Assessing, using, and replicating this work also requires access to documentation pertaining to participant characteristics, image acquisition and other scanning parameters, preprocessing and analysis procedures, as well as research materials including stimuli and custom code sets.

However, the importance of RDM to neuroimaging research extends beyond the need to save and organize multifarious sets of materials. The BOLD (Blood Oxygen Level Dependant) signal, which underlies the majority of functional MRI studies, has a complex origin^11^, is potentially confounded by a number of physical, physiological, and behavioral variables^12^, and requires careful interpretation^13^. The process of analyzing MRI data is also highly flexible, iterative, and statistically challenging, with decisions made at an early stage having significant downstream effects^14^. Operating system type, software versions, and even hardware architecture have also been shown to significantly influence analytical results^15^. Thus, extensive documentation and justification of data acquisition and analysis parameters is essential. Unfortunately, despite the publication of best practice guidelines^11^, many articles describing the results of projects involving MRI omit essential details related to experimental design, data acquisition, and analysis procedures^17,18^.

The rigor and reproducibility of neuroimaging research has been questioned due to a number of interrelated issues including reporting and publication biases in the scholarly literature^19^,^20^, low levels of statistical power^21^,^22^, the use of suboptimal design and analytical methods^23,24^ and the recent discovery of errors in widely used software tools^25^. Open data sharing has long been proposed as a way to address, if not necessarily resolve, these and other issues^26,27^. While sharing neuroimaging data does not directly address shortcomings in research design or analytical methodology within single studies, it does allow for their methods and results to be more fully examined and hopefully addressed in future work.

Early attempts such as the fMRI Data Center (fMRIDC) were met with skepticism and hampered by the demands involved in curating such large and complex datasets, an absence of community-wide requirements or incentives to make data available, and a lack of formalized standards about how neuroimaging data should be organized^28^. Though the Journal of Cognitive Neuroscience began to require the sharing of MRI data via the fMRIDC in 2000, this requirement was dropped by 2006 and data sharing requirements remain rare among neuroimaging journals. However, the view of the broader research community has since shifted considerably. Researchers increasingly appear to support the concept, if not the actual practice, of sharing their data^10,29^ and other data stakeholders including scholarly publishers^30^ and federal funding agencies^31^ have adopted a heterogeneous mix of data-related policies, mandates, and best practice recommendations. In parallel, a wide variety of tools and platforms have been developed to allow neuroimaging researchers to more easily manage and share MRI data.

Reflecting the iterative and flexible nature of how it is analyzed, MRI data is currently disseminated in a variety of forms. For example, tools like Neurosynth (http://neurosynth.org/) enable researchers to examine and compare peak activation coordinates reported in the neuroimaging literature while platforms such as Neurovault (https://neurovault.org/) and OpenfMRI (https://openfmri.org/) allow researchers to share the data in the form of unthresholded statistical maps and raw images respectively. Projects such as the Alzheimer’s Disease Neuroimaging Initiative (ADNI)^32^, the International Neuroimaging Data Sharing Initiative (INDI)^33^, and the Autism Brain Imaging Data Exchange (ABIDE)^34^ host large datasets related to clinical conditions and the National Institutes of Health sponsors the Connectome Coordination Facility (CCF) (https://www.humanconnectome.org/), which makes carefully collected, large-scale multimodal data available for reuse. The development of standardized organizational schemes such as the Brain Imaging Data Structure (BIDS)^35^ and tools for constructing and distributing analytical pipelines (e.g. LONI^36^, Nipype^37^, BIDS Apps^38^) enable researchers to not only share their data but also ensure that it can be navigated, assessed, and reproduced by others. However, while these developments have provided important infrastructure, addressing concerns related to rigor and reproducibility requires more than simply the adoption of new technologies. It also requires the refinement of a broad spectrum of behaviors and practices^39^.

Rigorous and reproducible science begins in the laboratory. For data to be effectively shared, evaluated, and re-used, it must first be effectively documented, organized, saved, and prepared. Such activities are encapsulated in the FAIR (<u>F</u>indable, <u>A</u>ccessible, <u>I</u>nteroperable, and <u>R</u>e-usable) Data Principles, which were developed by an international community of researchers, librarians, funders, and publishers as guidelines for enhancing the reusability of research data^40^. Though similar principles have been incorporated into recent neuroimaging-specific best practice recommendations^41^, the extent to which they translate into the day-to-day activities of active researchers remains unclear. Therefore, to inform efforts within both the neuroimaging and academic library communities to address rigor and reproducibility, we designed and distributed a survey examining RDM-related practices and perceptions among neuroimaging researchers working with MRI data.

## Methods

Our survey consisted of 74 multiple choice questions and an optional open response question. Questions focused on a range of data management-related topics, including types of data collected (including MRI data, non-MRI data, and related documentation), tools used to manage and analyze data, and the degree to which data management practices are standardized within the participant’s research group. The survey was distributed using the Qualtrics platform (http://www.qualtrics.com) between June and September, 2017. Before beginning the survey, participants were required to verify that they were at least 18 years of age and gave their informed consent to participate. Participants were able to skip questions while proceeding through the survey. All study procedures were approved by the institutional review board of Carnegie Mellon University (Study 2017_00000129) and all research was performed in accordance with relevant guidelines and regulations. Data were analyzed using JASP^42^.

### Survey Design

The process of survey design drew upon expertise from both the academic library and neuroimaging communities. The structure of the survey, which generally follows the trajectory of a typical MRI project, drew from the research data lifecycle - a model that has been widely adopted by data support providers in academic libraries to organize activities related to the management of data over the course of a research project^43^. Building on similarly-structured tools, such as the data curation profiles^44^, survey questions were developed in consultation with researchers actively working in the field to ensure that each question was tailored to the specific terminology, practices, and tools currently employed by the MRI research community. Whenever possible, we referred to RDM-related concepts and activities using language that would be familiar to neuroimaging researchers. We also avoided using terms like “metadata”, which carry potentially different meanings within the context of MRI research (e.g. information included in the header of image files) and research data management more broadly (e.g. standards such as Dublin Core or any documentation that provides “data about data”).

For questions referencing data management maturity, we drew upon the capability maturity model framework^45^ which describes activities based on their degree of definition, standardization, and optimization. For the purposes of this study, RDM maturity was defined as the extent to which data management practices are clearly defined, implemented, and (if applicable) optimized. Introductory text also clarified that the survey was not designed to judge researchers who have different styles of data management or whose practices exhibit different levels of sophistication. Though capability models specific to the management of scientific data have been developed^46^, to our knowledge this is the first survey to apply this framework as a means to collect quantitative maturity ratings from the research community itself.

Because we believed that participants would come to our survey with different perspectives on RDM-related topics and terms, each section of the survey was preceded by a brief description of the specific activities and practices covered in that section as well as an operational definition of data management maturity. The final section of the survey covered emerging publication practices such as preregistering studies, disseminating preprints, and publishing open access articles. Though such practices are do not technically fall within the bounds of RDM, they represent overlapping efforts to address the transparency, rigor, and reproducibility of the research process.

### Distribution and Filtering Criteria

Recruitment emails were sent to the directors of neuroimaging centers and other MRI facilities affiliated with universities and other research institutions. The survey was also advertised through social media and via psychology, neuroscience, and neuroimaging-related discussion groups and mailing lists (e.g. the SPM and FSL mailing lists, the PsychMap Facebook group, the BrainHack Slack channel). Though we initially intended to collaborate with neuroimaging-focused scientific societies on the distribution of our survey, we were unable to make arrangements to do so before the close of data collection.

In order to capture a broad view of data management-related practices and perceptions inclusion criteria were only that potential participants be an active researcher using MRI, that they were over the age of 18, and that they consented to participate in our study. Data from participants who did not meet these criteria or who completed less than a single section of the survey were excluded from subsequent analyses.

### Data Availability

The survey instrument (including consent documentation and instructions)47 and resulting dataset (excluding personally identifying information)48 are both available via figshare.

## Results

### Participant Characteristics

A total of 144 participants from 11 countries and 69 institutions participated in this study. The majority of participants were from the United States (72.31%), the United Kingdom (10.00%), and Canada (6.92%). As shown in Table 1, participants were affiliated with a variety of research disciplines with the most common being cognitive neuroscience. Participants consisted of a mix of trainees (e.g. graduate students, postdoctoral fellows) and faculty (e.g. associate, assistant, and full professors).

**Table 1.** Characteristics of study participants. A total of 144 neuroimaging researchers participated in this study, though not every participant gave a response for every question. Participants were split between **A.** trainees and faculty and **B.** cognitive neuroscience and other research areas. All values listed are percentages.

| A. Participant title or role |  | B. Participant research area |  |
| --- | --- | --- | --- |
| Title | Percent | Research Area | Percent |
| Graduate Student | 24.31 | Cognitive Neuroscience | 55.56 |
| Assistant Professor | 21.53 | Clinical Neuroscience | 15.97 |
| Postdoctoral Fellow | 21.53 | Developmental Neuroscience | 5.56 |
| Associate Professor | 10.42 | Social Neuroscience | 5.56 |
| Professor | 9.03 | Behavioral Neuroscience | 2.08 |
| Research Associate/Scientist | 6.94 | Computational Neuroscience | 2.08 |
| Research Assistant | 2.08 | MRI Methods | 2.08 |
| Research Technician | 1.4 | Bio/Neuroinformatics | 1.39 |
| Staff Scientist | 1.4 | Sensory Systems Neuroscience | 1.39 |
| Other | 1.4 | Affective Neuroscience | 1.39 |
| *N* = 130 |  | Other | 6.9 |
|  |  | *N* = 144 |  |

The majority of participants (64.06%, *N* = 128) indicated that they receive funding from the National Institutes of Health (NIH). Other common sources of funding include private foundations (12.5%), the National Science Foundation (NSF) (11.72%), internal grants (including startup funds) (11.72%), the Department of Defense (DOD) (3.90%), and international funding bodies (19.53%). Though most participants (53.47%, *N* = 144) indicated that they generate documentation about how data is to be collected, organized, and secured over the course of a project, the heterogeneous data policies of these funding bodies makes it highly unlikely that this refers exclusively to a data management plan (DMP).

Participants indicated that they received their training about how to collect and analyze neuroimaging data from a variety of sources, including individuals within their own lab (e.g. other students, post-docs) (57.64%, *N* = 144), online resources and documentation (e.g. self-taught) (51.39%), individuals outside their lab (e.g. other people in their department) (35.42%), and formal courses at their institution (20.14%) or another institution (27.08%). A relatively small percentage of participants indicated that they had taken advantage of local university services related to data management (27.8%) or scholarly publishing (14.6%), though engagement with services related to technical infrastructure was considerably higher (45.1%). Instead, the majority indicated that such services were unavailable, that they were unsure about their existence, or that they were aware of their existence but had not taken advantage of them.

### RDM Maturity Ratings

As shown in Figure 1, participants rated the overall maturity of their data management practices during both the data collection [*t*(128) = 6.349, *p* < 0.001] and data analysis [*t*(116) = 7.403, *p* < 0.001] phases of a project as significantly higher than those of the field as a whole. A similar trend was observed for the data sharing phase, but the comparison did not reach statistical significance [*t*(115) = 1.677, *p* < 0.096]. Average maturity ratings for the data sharing phase were significantly lower than those of the other phases for both individual practices [*F*(2, 226) = 70.61, p < 0.001] and the field as a whole *F*(2, 226) = 34.44, p < 0.001].

**Figure 1.**
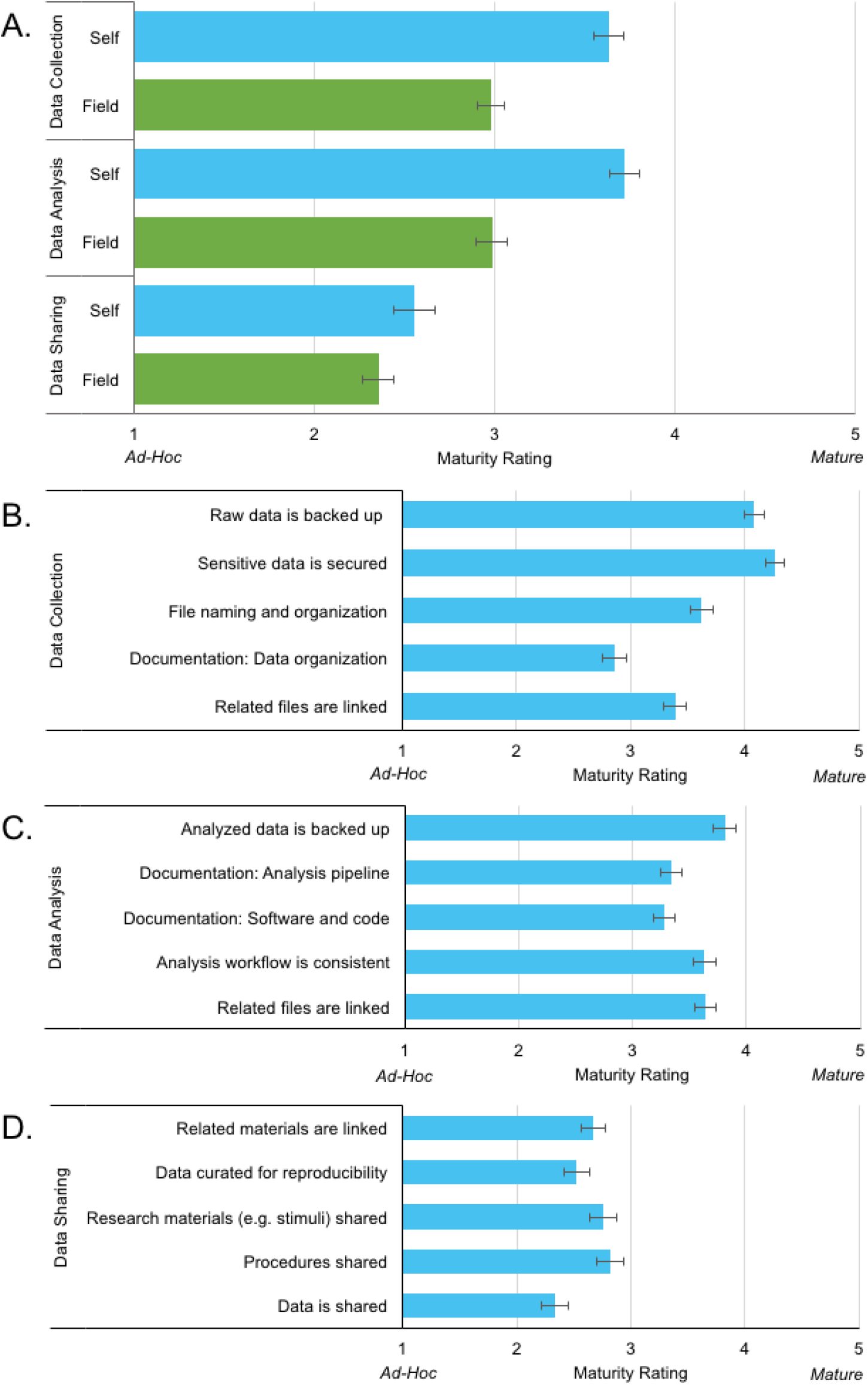
Average RDM maturity ratings between (A) and within (B-D) three phases of an MRI research project. **A.** Participants rated their own RDM practices as significantly more mature than those of the field as a whole during the data collection and analysis phases. Ratings of both individual and field maturity were significantly lower during the data sharing phase than during data collection and analysis. [Data collection: *N* =131 (individual), 130 (field), data analysis: *N* = 118 (individual/field), data sharing: 116 (individual/field)]. Ratings of individual activities within each phase reflected a similar trend. **B.** Practices related to the backup of raw data and securing of sensitive data were rated as highly mature during the data collection phase while the documentation of file organization schemes (such as through a lab notebook or data dictionary) received the lowest rating. [*N* = 132] **C.** Similarly, during the data analysis phase, the backup of analyzed data received the highest rating, while the documentation of decisions related to analytical pipelines and the use of computational tools received the lowest. [*N* =120] **D.** Activities described in the data sharing phase received lower ratings than those in previous phases. [*N* = 116]

Ratings of individual practices within each phase followed a similar pattern. Overall, ratings for individual practices during the data sharing phase were substantially lower than those during the data collection and analysis phases. Maturity ratings were highest for practices involved in ensuring the security of sensitive data and backing up data and lowest for those involved making data available to researchers outside of their research group. This focus on practical concerns was also evident when participants were asked about what motivates and limits their RDM practices. As shown in Table 2, participants reported that they were primarily motivated by a desire to prevent the loss of data, ensure everyone in their research group has access to data, and a desire to foster openness and reproducibility and limited by time and a lack of available training and best practices.

**Table 2.** Factors that limit and motivate RDM during the data collection, analysis, and sharing/publishing phases of a research project. All values listed are percentages, more than one response could be selected. In terms of limits, “Other” responses included changes in personnel, differences in expertise within a lab, differences in preferences between lab members, lack of top-down leadership, and concerns about future cost. For motivations, “other” responses included ensuring continuity following personnel changes, keeping track of analyses, error prevention, and maximizing efficiency. [Data collection: *N* = 125 (limits/motivations), Data analysis: *N* = 115 (limits), 120 (motivations), Data sharing: *N* =112 (limits/motivations)].

| RDM limits and motivations |  | Data Collection | Data Analysis | Data Sharing |
| --- | --- | --- | --- | --- |
| **Limits** | The amount of time it takes | 69.60 | 71.30 | 79.46 |
|  | Lack of best practices | 43.20 | 48.70 | 49.11 |
|  | Lack of incentives | 36.80 | 32.18 | 37.50 |
|  | Lack of knowledge/training | 32.80 | 40.87 | 41.07 |
|  | The financial cost | 17.60 | 8.70 | 22.32 |
|  | Other | 7.20 | 6.09 | 5.36 |
| **Motivations** | Prevent loss of data | 100.00 | 85.83 | 78.57 |
|  | Ensure access for collaborators | 76.80 | 73.33 | 70.53 |
|  | Openness and reproducibility | 63.20 | 64.17 | 66.96 |
|  | Institutional data policy | 52.00 | 39.17 | 47.32 |
|  | Publisher/Funder Mandates | 35.20 | 28.33 | 41.96 |
|  | Availability of tools | 12.00 | 9.17 | 8.93 |
|  | Other | 3.20 | 3.3 | 0.0 |

### Data Collection Practices

For the purposes of this survey, “data collection” was defined as activities starting with the collection of neuroimaging data at a scanning facility and continuing through to the organization and storage of data within the participant’s laboratory. Questions in this section of the survey dealt primarily with the types of data collected as well as procedures for moving, saving, and organizing raw data.

As expected, participants indicated that they collect and manage a wide range of research materials over the course of an MRI project. As shown in Table 3, this includes multiple types of MRI images, additional “non-MRI” data, and a variety of documentation, code, and other research materials related to data collection and analysis. When it comes to moving data from the scanning facility, the majority of participants indicated that they use a server to transfer their MRI data (82.58%, *N* = 132) and a hard drive for non-MRI data (55.30%). Once data is in the lab, participants indicated that it is primarily organized using standardized file structures (70.45%, *N* = 132) and file names (67.42%). Less common were the use of formal lab notebooks (47.73%), databases (28.79%), or the admission that procedures are generally not documented (17.42%). The majority of participants indicated that practices related to data organization were consistent within their lab or research group. However, trainees were significantly less likely to endorse this consistency than faculty members [*X*^2^(2, *N* = 132) = 13.49, *p* < 0.01]. A similar trend was evident when participants were asked about backup procedures. The majority of participants (73.5%, *N* = 132) indicated that their scanning facility maintains backups of MRI data and that they themselves backup data using a wide variety of means including servers operated by their lab (41.67%, *N* =132) or institution (37.88%), manual (30.30%) and automatic (21.97%) backups of local machines, and external hard drives (25.76%). Though the majority of participants (54.2%) indicated that backup procedures were consistent within their lab or research group, trainees were less likely to endorse this than faculty [*X*^2^(2, *N* = 131) = 7.28, *p* < 0.05].

**Table 3.** Types of data collected fell into three categories: MRI data, non-MRI data, and study information. All values listed are percentages, multiple data types could be selected. **A.** For MRI data, common “other” responses included spectroscopy, diffusion, blood flow, and MRS. **B.** For non-MRI data, common “Other responses included motion tracking, neurophysiology measures, and hormones (saliva). **C.** For study information, common “other” responses included scanner quality assurance data, information about the scanner itself, and consent forms.

| A. MRI Data |  | B. Non-MRI data |  | C. Study Information |  |
| --- | --- | --- | --- | --- | --- |
| Data | Percent | Data | Percent | Data | Percent |
| Anatomical | 99.24 | Demographics | 97.0 | Acquisition parameters | 97.7 |
| Task-related | 98.54 | Behavioral data | 95.5 | Task information | 97.7 |
| Resting state | 80.30 | Questionnaires | 88.6 | Stimuli | 91.7 |
| Field map | 64.46 | Clinical data | 60.6 | Session information | 90.2 |
| Diffusion | 59.84 | Physiological data | 43.2 | Code (presentation) | 82.6 |
| Other | 11.36 | Genetic data | 30.3 | Code (data collection) | 71.2 |
|  |  | Other imaging | 25.0 | Other | 7.6 |
|  |  | Other | 3.8 |  | *N* = 132 |

### Data Analysis Practices

For the purposes of this survey, “data analysis” was defined as activities starting with preprocessing (co-registration, motion correction, etc) of MRI data and proceeding through first and second level analyses. Questions in this section of the survey dealt primarily with the use of software tools and the documentation of analytical decisions and parameters. Overall, participants indicated that they use a wide variety of tools to analyze their MRI and non-MRI data. While there are several commonly used tools, as shown in Table 4, there is also a long tail of tools used by a relatively small number.

**Table 4.**
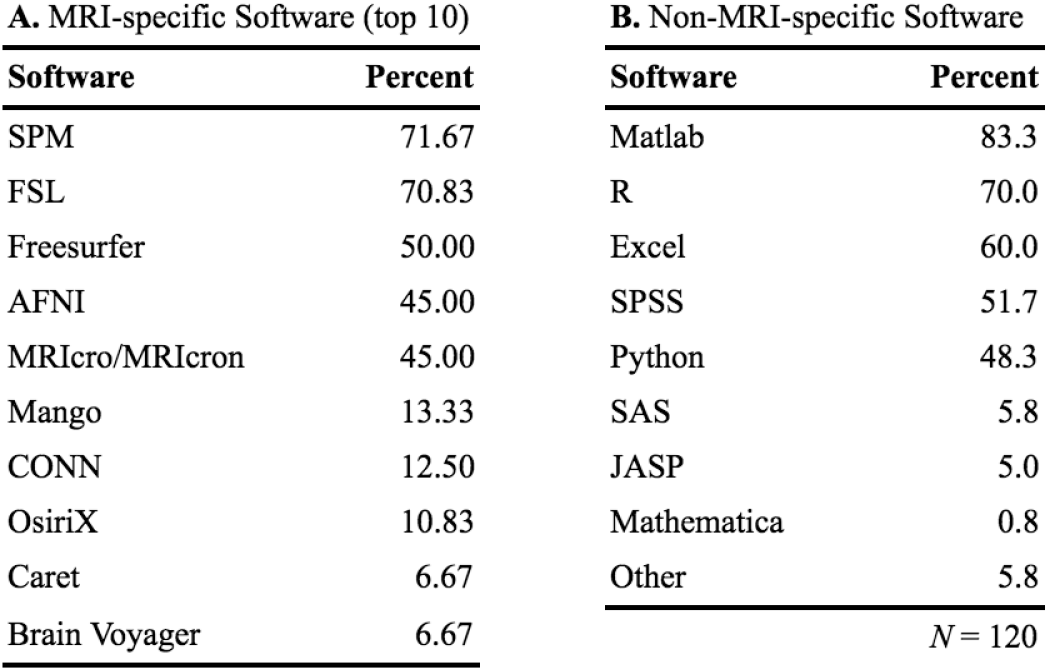
Software used for analysis: **A.** MRI-specific software (top 10 most popular) and B. non-MRI-specific software. All values listed are percentages, multiple software tools could be selected. “Other” MRI-specific software included Nipype (5.00%), custom code (4.17%), ANTS (4.17%), FMRIprep (2.50%), NiPy (1.17%), ITK-SNAP (1.17%), Connectome Workbench (1.17%), MRIQC (1.17%), CIVET, C-PAC, DPARSF, GIFT, ExploreDTI, CAT, SPHARM, TBSS, fidl, PLS, SamSrF, Vistasoft, and MedIRNIA. “Other” non-MRI-specific software included Acknowledge, CIGAL, Fscan, Data Desk, Mplus, Octave, Stan, and Bash.

Only 13.33% of participants (*N* = 120) indicated that they process data from each subject individually using a GUI. Instead, the majority of participants indicated that their preprocessing is scripted using their own scripts (64.17%), scripts adapted from others (58.33%), or scripts written by others without adaptation (15.0%). Many participants indicated that everyone in their lab or research group uses the same tools to analyze MRI data (40% indicated that everyone in their group also uses the same version of software tools, 25% indicated that group members use different versions). Trainees were again significantly less likely to endorse this than faculty [*X*^2^(3, *N* = 120) = 25.4, *p* < 0.001]. Analysis of non-MRI software tools yielded similar results, though only 43.3% (23.3% same version, 20% different versions) of participants indicated that the application of these tools was consistent. Again, differences between trainees and faculty were statistically significant [*X*^2^(2, *N* = 120) = 14.90, *p* < 0.01]

Participants indicated that they generally document their activities (including quality checks, pre-processing parameters, and the results of first/higher level analysis) using a word processing program (e.g. Evernote, Microsoft Word) (56.67%, *N* = 120), readme files (42.5%), and, to a lesser extent, version control systems (e.g. Git) (25.83%), electronic lab notebooks (e.g. Jupyter) (19.17%), active data management plans (4.17%), and lab management tools (e.g. LabGuru, Open Science Framework) (2.5%). Unfortunately, 10.83% of participants indicated that they do not document their activities in any systematic way. The majority of participants (74.6%) acknowledged that not everyone uses the same system for documenting their activities and differences between trainees and faculty were again statistically significant [*X*^2^(2, *N* =118)= 17.55, *p* < 0.001].

When asked if another researcher could recreate their work using only its documentation, 59.2% of participants (*N* = 120) indicated that they would be able to recreate both their preprocessing and analysis steps, 11.5% indicated that other researchers would be able to recreate one or the other, and 19.1% were either unsure or believed that they would need to be present.

### Data sharing Practices

For the purpose of this survey, “data sharing” was defined as activities involving the dissemination of conclusions drawn from neuroimaging data as well as the sharing of the underlying data itself through a general or discipline-specific repository. This definition was made intentionally broad in order to capture the fact that MRI data is currently made available in a variety of forms, ranging from coordinate tables published in the neuroimaging literature to raw and preprocessed datasets deposited in repositories alongside relevant code, stimuli, and documentation. Questions in this section of the survey dealt primarily with the means and motivations for making data, code, and other materials available to other researchers.

As shown in Table 5, participants generally indicated that they were motivated to share data by a desire to foster research transparency and reproducibility rather than by professional incentives or the need to fulfill mandates. Whhen asked about reasons they may not be able to share their data, the most common responses were that it may contain additional findings to be discovered or published and that it contains confidential or sensitive information. Half (50.0%, *N* =116) of participants indicated that they had not been required to share data or submit a data availability statement when publishing a journal article. However, significantly more faculty (50.88%) indicated that they had encountered such a requirement than trainees (28.81%) [*X*^2^(3, *N* =116)= 10.52, *p* < 0.05]. Similarly, while the majority of participants indicated that they have not requested data from an author of a journal article (56.03%) or received such a request themselves (55.17%), significantly more faculty reported receiving a request for their data than trainees [*X*^2^(2, *N* =116) = 21.62, *p* < 0.001].

**Table 5.** Reasons why data can and cannot be shared. All values listed are percentages, more than one reason could be selected. Other reasons given include: Consent (5), laziness, afraid of mishandling, projects that are haphazard.

| A. Reasons for sharing data |  | B. Reasons for not sharing data |  |
| --- | --- | --- | --- |
| Reason | Percent | Reason | Percent |
| Transparency and openness | 55.14 | Data contains additional findings to publish | 50.43 |
| To enable reuse/reproducibility | 55.14 | Data contains sensitive information | 30.43 |
| To communicate my results | 49.53 | It would take too much time | 25.22 |
| To allow others to check my work | 47.66 | Supervisor doesn't wish to share | 16.52 |
| Incentives (authorship, citations) | 21.50 | Format of data make sharing difficult | 15.62 |
| Mandated by funder/publisher | 20.56 | Don't know how | 14.78 |
| Establish intellectual property | 1.87 | Data is proprietary, subject to IP | 1.74 |
| Other | 4.67 | Other | 7.82 |
| NA | 28.04 | Can share, but require citation | 29.57 |
|  |  | Can share, but require authorship | 9.57 |
|  |  | N = 115 |  |

Participants indicated that a large number of their research materials should be preserved over the long term (see Table 6) and generally reporting saving materials for eight years or more (29.31% maintained so that is always accessible, 40.51% saved in formats that may become obsolete). When asked if another researcher could recreate their work using only its description in a publication or scholarly report, 64.38% (*N* = 115) indicated that they would be able to recreate both their preprocessing and analysis steps, 9.57% indicated that other researchers would be able to recreate one or the other, and 26.09% were either unsure or believed that they would need to be present.

**Table 6.**
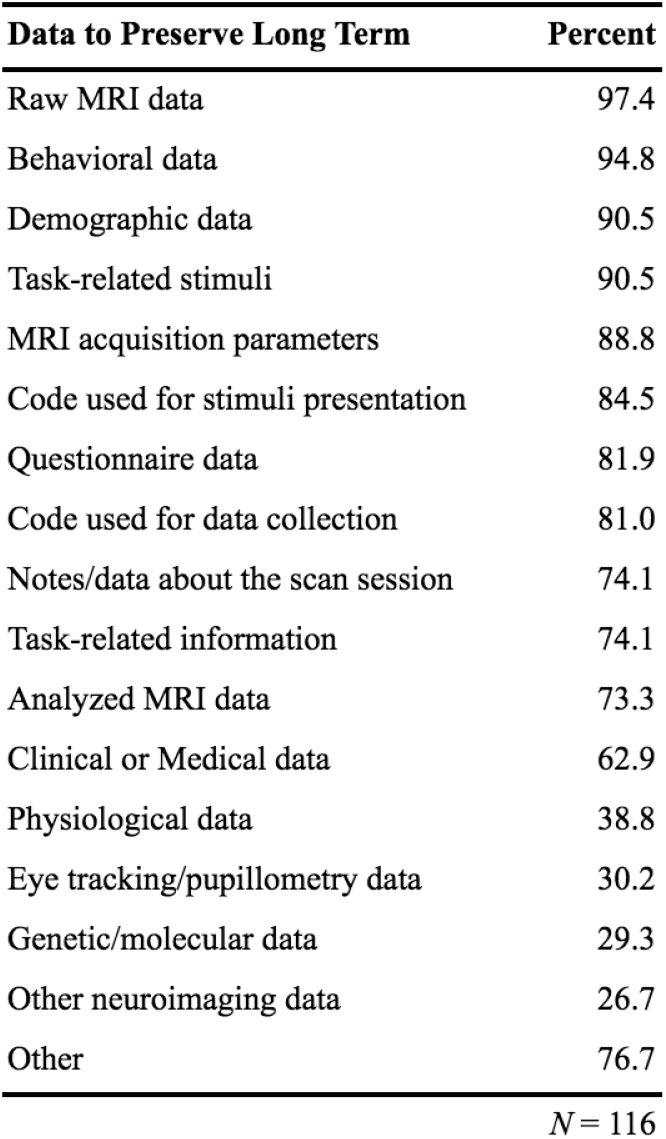
Important parts of data to preserve long term. All values listed are percentages, multiple data types could be selected. Overall, researchers want to preserve nearly all data long term. Other data types indicated to preserve include: code for analysis (3) and hormone information.

### Emerging Research Practices

The majority of participants (56.64%, *N* =113) indicated that they currently regard data as a “first class” research product, meaning a product that should be assessed, valued, and considered as part of application and promotion decisions in the same way as a journal article. As shown in Figure 2, this is broadly indicative of that fact that the MRI research community is currently at a point of transition. While only a small percentage of researchers indicated that they have adopted emerging research practices such as pre-registering their studies, conducting replications, publishing preprints, or publishing research products such as code, datasets, and grant proposals, a substantial percentage indicated that they plan to in the future.

**Figure 2.**
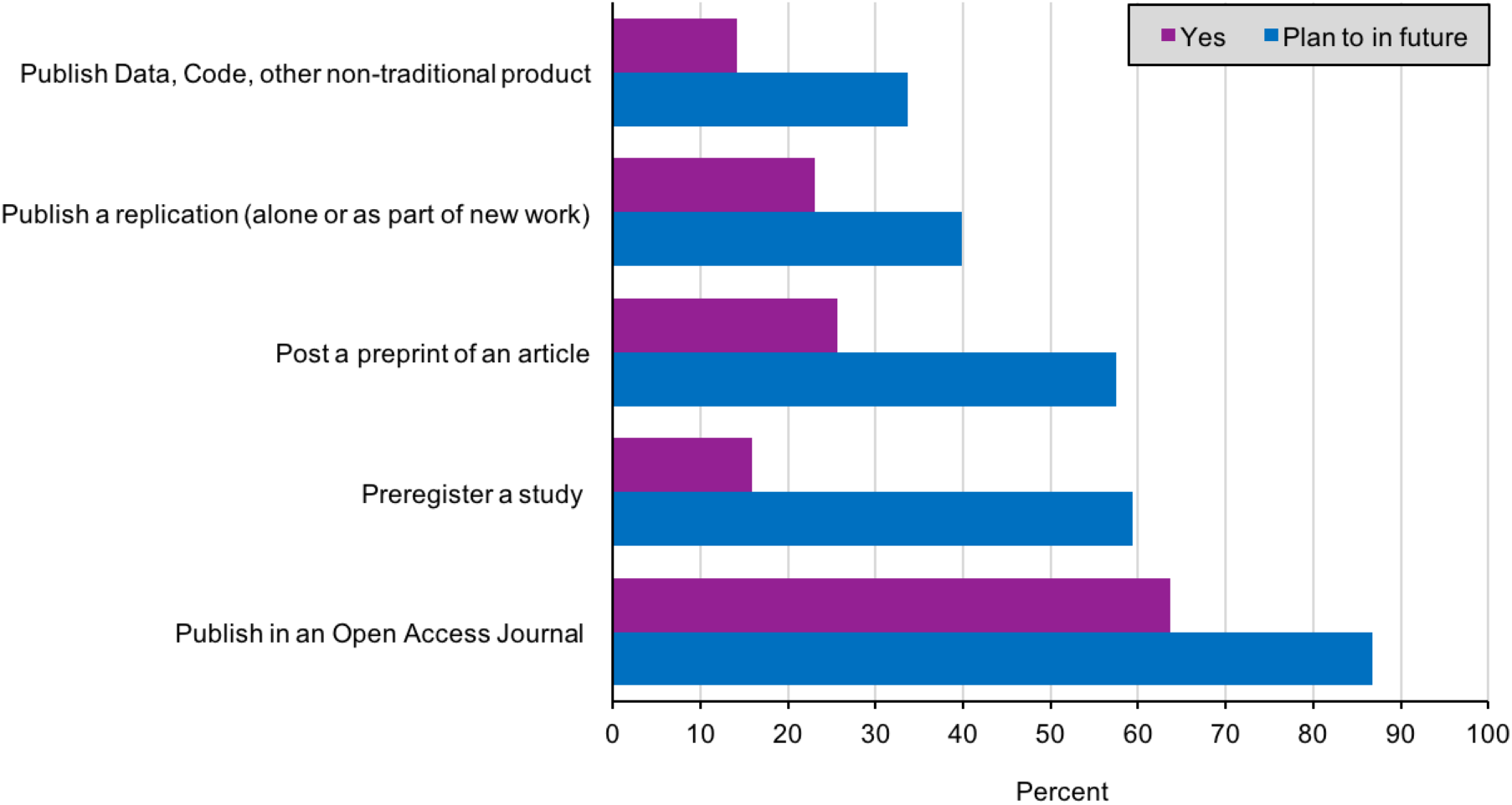
Adoption of emerging research practices among neuroimaging researchers. [*N* = 100]

## Discussion

In order to inform efforts within the neuroimaging and academic library communities to address issues of rigor and reproducibility, we surveyed the RDM-related practices and perceptions of researchers who use magnetic resonance imaging (MRI) to study human neuroscience. Overall, our results highlight the considerable challenges involved in properly managing and sharing neuroimaging data - the data is acquired in multifarious forms, transformed and analyzed using a wide variety of tools, and documented inconsistently. Our results also demonstrate that neuroimaging researchers generally receive informal training in data-related practices, have little interaction with institutional data support services, and presently encounter few expectations from data stakeholders such as scholarly publishers and research funding bodies.

Neuroimaging is not unique in facing challenges related to rigor and reproducibility. Issues such as publication bias^49,50^ and low statistical power^51,52^ have been discussed in the behavioral and biomedical sciences for decades and data stakeholders including scholarly publishers and federal funding agencies have instituted a range of reproducibility-related policies stipulating how the data underlying published work should be managed and shared. For example, while mandates requiring authors to share the data underlying publications have been shown to increase the degree to which data is made available^53^, only a minority of biomedical journals have such requirements and even fewer provide specific guidance as to how to make data available and reusable^54^. Federal funding bodies generally exercise their RDM-related policies by requiring that a data management plan (DMP), which outlines how data is going to be collected, organized, preserved, and shared, be submitted as part of any grant proposal^31^. The efficacy of DMPs in affecting how researchers actually manage and share their data in practice is unclear^55^ and it is notable that the NIH, which was the most prevalent funder in our sample, does not currently have a DMP requirement. However, as evidenced by the imminent pilot of an NIH Data Commons and the recent controversy related to the reclassification of behavioral and imaging studies as clinical trials^56^, funder data policies will likely soon begin affecting neuroimaging researchers.

Neuroimaging is also not unique in how it has addressed challenges related to rigor and reproducibility. Communities throughout neuroscience working with specific methodologies (e.g. neurophysiology^57^, cell morphology^58^) or data from specific model systems (e.g. C. elegans^59^) have begun to develop standards, tools, and community norms to facilitate the management, curation, and sharing of data and other research materials. Like complementary initiatives across other disciplines^60^, these grassroots efforts have the potential to ensure that effective RDM practices are incorporated throughout the course of a research project rather than simply deployed at discrete points in response to mandates from a funder or publisher. By assessing current RDM practices in neuroimaging, the present study adds crucial context to efforts aimed at advancing rigor and reproducibility.

By collecting and analyzing quantitative ratings of RDM maturity, which we operationalized as the degree to which data-related procedures are defined and implemented, we were able to quantify how active neuroimaging researchers perceive their own practices and the practices of the field as a whole. There are several interpretations of our observation that participants generally rated their own practices as more mature than those of the field as a whole. It is possible that this observation reflects the well-known phenomenon of participants rating themselves as better than average across a wide range of personal characteristics61. Given that this study was primarily disseminated via social media and through a number of scholarly discussion groups where there is a great deal of discussion related to research methodology, open science, and reproducibility, it is also possible that our sample was indeed biased in favor of participants who incorporate RDM into their work to a greater degree than average. Our finding that maturity ratings were significantly lower for the data sharing/publishing phase is in line with the lack of existing data sharing requirements, the propensity of researchers to share data via personal communication, and the centering of data sharing as a way to address issues of rigor and reproducibility. However, it is also at odds with the fact that our results indicate that there is ample room for improving RDM practices during the data collection and analysis phases.

By asking participants about their RDM-related activities using language and terminology familiar to them, we were able to construct a comprehensive picture of how neuroimaging researchers handle their data over the course of a project. Given the preponderance of informal methodological training in neuroimaging and results of previous studies examining methods reporting in the extant literature^19^,^20^, we expected that participants would report applying a wide variety of practices and tools to their data. Our results bore this out and also revealed that trainees and faculty members had significantly different perspectives on the degree to which backup procedures, data structures, analytical tools, and documentation practices are consistent within their lab or research group. While our methods do not allow us to speculate about the cause of these differences, their ubiquity indicates that there is not an optimal amount of communication about the importance of RDM even within individual research groups or projects.

The spread of increasingly high resolution imaging hardware, the rapid evolution of experimental approaches and analytical techniques, and the community development of user-friendly software tools have enabled neuroimaging researchers using MRI to make significant contributions across the behavioral and biomedical sciences. However, our results demonstrate there is an outstanding need for training related to research data management. We hope that the results of this assessment can inform the development of materials and training for RDM practices in neuroimaging and the creation of tools to support these practices. This issue is also not unique to neuroimaging. In a recent survey, PI’s from across the National Science Foundation (NSF)’s biological sciences directorate listed training on data management as one of their foremost unmet needs^62^. Though topics related to RDM are often included in undergraduate and graduate level coursework, many educators report that they are not covered thoroughly due to a lack of time, expertise, and guidance^63^. These trends highlight the need for greater collaboration between researchers who possess expertise in collecting, analyzing, and evaluating data and support providers in academic libraries who have expertise in its management and sharing. Because our results demonstrate that RDM activities among neuroimaging researchers are, at least at present, generally motivated by immediate considerations such as ensuring data is not lost and ensuring that it is accessible to the members of a particular research group, a potentially fruitful direction for such a collaboration could be the development of training materials that provide actionable information about how data could be effectively documented, organized, and saved throughout the course of a research project and also illustrate how such activities are an important component of addressing broader concerns related to rigor and reproducibility. As academic libraries transition to an era of digital resources and growing expertise in data management, data science, and open science, librarians and library-based service providers are well positioned to become trusted consultants and active collaborators in labs and scanning centers and to provide training to students and researchers on RDM practices. Several global initiatives such as the Research Data Alliance (RDA) and Force 11 already bring together researchers, librarians, and other data stakeholder groups to address complex issues related to research data. Increased collaboration between such groups in the design and implementation of neuroimaging-specific resources could be quite fruitful in the future. In particular, library professionals could specifically address how research data fits into the broader scholarly communications ecosystem and apply their expertise in both RDM and scholarly publishing to issues facing the neuroimaging in conjunction with the technical solutions being developed.

Though the present study offers unique insight into the data management practices of neuroimaging researchers, these results should not be interpreted as a criticism or singling-out of the field. Follow-up research will explore RDM practices and perceptions in cognate research areas such as psychology and biomedical science and it is likely that many of the same trends-including informal education in effective RDM practices, inconsistency even within the same research group, and slow adoption of open science tools and practices will be observed. Follow-up research will also explore researchers’ expertise with different data management practices as well as the degree to which researchers are willing and able to change their practices when presented with alternative approaches and evolving requirements from funders and publishers.

## Acknowledgements

This work was partially funded by a Berkman Faculty Development Grant awarded to A.V. by Carnegie Mellon University. While conducting the work described in this publication, J.B. was funded as both a CLIR Software Curation Fellow (Alfred P. Sloan Foundation #G-2015-14112) and an RDA Data Share Fellow (Alfred P. Sloan Foundation #G-2014-13746, National Science Foundation NSF ACI #1349002). The authors would like to thank Yael Isler and John Pyles for their helpful comments throughout the research process and J.B. Poline and Russ Poldrack for their suggestions related to the survey instrument.

## Author contributions statement

J.B. and A.V. jointly conceived the study, designed the survey, analyzed the results, and wrote the manuscript.

